# Engineering *Escherichia coli* to produce aromatic chemicals from ethylene glycol

**DOI:** 10.1101/2023.01.08.523183

**Authors:** Smaranika Panda, Jie Fu J Zhou, Michelle Feigis, Emma Harrison, Xiaoqiang Ma, Vincent Fung Kin Yuen, Radhakrishnan Mahadevan, Kang Zhou

**Affiliations:** Department of Chemical and Biomolecular Engineering, National University of Singapore; Department of Chemical Engineering and Applied Chemistry, University of Toronto

**Keywords:** Ethylene glycol, Aromatic chemicals, *Escherichia coli*, Metabolic engineering, L-tyrosine, Polyethylene terephthalate (PET)

## Abstract

Microbial overproduction of aromatic chemicals has gained considerable industrial interest and various metabolic engineering approaches have been employed in recent years to address the associated challenges. So far, most studies have used sugars (mostly glucose) or glycerol as the primary carbon source. In this study, we used ethylene glycol (EG) as the main carbon substrate. EG could be obtained from the degradation of plastic and cellulosic wastes. As a proof of concept, *Escherichia coli* was engineered to transform EG into L-tyrosine, a valuable aromatic amino acid. Under the best fermentation condition, the strain produced 2 g/L L-tyrosine from 10 g/L EG at approximately 50% of the theoretical yield, outperforming glucose (the most common sugar feedstock) in the same experimental conditions. To prove the concept that EG can be converted into different aromatic chemicals, *E. coli* was further engineered with a similar approach to synthesize other valuable aromatic chemicals, L-phenylalanine and *p*-coumaric acid. Finally, waste polyethylene terephthalate (PET) bottles were degraded using acid hydrolysis and the resulting monomer EG was transformed into L-tyrosine using the engineered *E. coli*, yielding a comparable titer to that obtained using commercial EG. The strains developed in this study should be valuable to the community for producing valuable aromatics from EG.

## 1. Introduction

Aromatic amino acids and their derivatives have important applications in the chemical, food, and pharmaceutical sectors (Huccetogullari et al., 2019). Among them, L-tyrosine has received considerable attention, as it is a key precursor to 1-3,4-dihydroxyphenylalanine (L-DOPA, a frontline drug for treating Parkinson’s disease) and many other natural products including flavonoids and alkaloids (Hinz et al., 2011; Kim et al., 2018).

In nature, aromatic amino acids are synthesized through the shikimate pathway, which begins with the condensation of phosphoenolpyruvate (PEP, an intermediate of glycolysis) and erythrose-4-phosphate (E4P, an intermediate of pentose phosphate pathway) to form 3-deoxy-D-arabino-heptulosonate (DAHP), which was catalyzed by the DAHP synthase (encoded by *aroG* in *Escherichia coli*). Through six reactions, DAHP is condensed with another molecule of PEP and transformed into chorismate, a crucial precursor to the three aromatic proteinogenic amino acids (**Fig. 1**). To synthesize L-tyrosine, chorismate is rearranged and oxidatively decarboxylated to form 4-hydroxyphenylpyruvate (HPP). One protein catalyzes this transformation in *E. coli* and is encoded by *tyrA*. HPP can be converted into L-tyrosine through transamination using L-glutamate as the amino group donor (**Fig. 1**). AroG and TyrA were found to catalyze rate-limiting steps in the pathway.

**Fig. 1.**
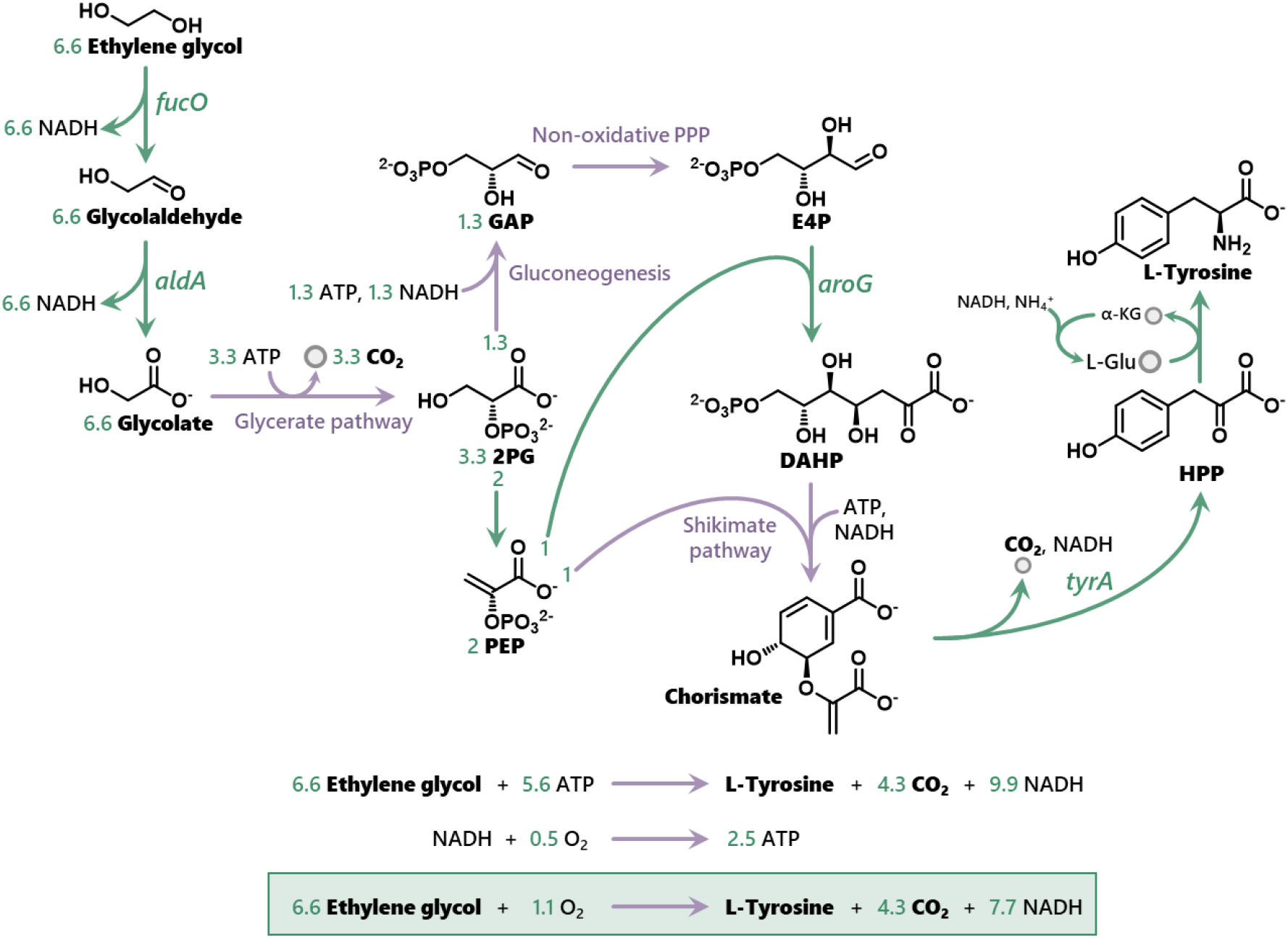
The metabolic pathway to synthesize L-tyrosine from ethylene glycol. 2PG: 2-phosphoglycerate. PEP: phosphoenolpyruvate. GAP: glyceraldehyde 3-phosphate. E4P: erythrose 4-phosphate. DAHP: 3-deoxy-D-arabino-heptulosonate 7-phosphate. HPP: 4-hydroxyphenylpyruvate. L-Glu: L-glutamate. α-KG: α-ketoglutarate. *fucO* encodes L-1,2-propanediol oxidoreductase. *aldA* encodes an aldehyde dehydrogenase. *aroG* encodes DAHP synthase. *tyrA* encodes chorismate mutase/prephenate dehydratase.

Many studies have been done to improve the microbial overproduction of aromatic chemicals in *Escherichia coli* (Averesch and Kromer, 2018; Huccetogullari et al., 2019; Kim et al., 2018; Lutke-Eversloh and Stephanopoulos, 2007; Ma et al., 2020; Rodriguez et al., 2014; Yang et al., 2018). So far, all the employed metabolic engineering approaches relied mainly on sugars (especially glucose) or glycerol as the primary carbon source. The high production cost, which is primarily due to the use of sugar-based feedstock and the relatively low product yield, is one of the major bottlenecks in commercializing fermentative production of these chemicals (Rosales-Calderon and Arantes, 2019). These facts motivated us to evaluate non-conventional substrates that may have lower substrate costs and lead to higher product yield.

One promising substrate candidate is ethylene glycol (EG), which could be derived from polyethylene terephthalate (PET) wastes through chemical (Carta et al., 2003; Stanica-Ezeanu and Matei, 2021) or enzymatic hydrolysis (Knott et al., 2020; Lu et al., 2022; Maurya et al., 2020). It is more difficult/expensive to recycle the EG fraction of the PET hydrolysate than the terephthalate fraction, because of EG’s high solubility in water and high boiling point (Tournier et al., 2020). EG could also be obtained from cellulosic wastes via a one-pot heterogeneous catalytic process that involves hydrolysis, retro-aldol condensation, and hydrogenation (Hamdy et al., 2017; Ji et al., 2009; Wang and Zhang, 2013; Yan et al., 2008; Yu et al., 2018). The process avoids the use of hydrolyzing enzymes but has not been commercialized mainly due to the high purification cost of EG. A few recent studies have also established the electrochemical reduction of CO_2_ into EG (Chen et al., 2021; Tamura et al., 2015). Theoretically, the crude EG solution produced by the above processes could be used as carbon substrate to fuel microbial fermentation, by-passing the EG purification challenge. We have also shown that EG utilization pathway is orthogonal to native metabolism (Pandit et al., 2021) of the cell thereby enabling the ease of engineering production pathways from EG.

Recently, we have improved growth of an engineered *E. coli* strain on EG through genetic engineering and medium optimization (Panda et al., 2021), and demonstrated production of glycolate (Pandit et al., 2021). So far, microbial conversion of EG into value-added chemicals other than glycolate has not been reported.

Here, we have engineered *E. coli* to synthesize L-tyrosine using EG as the major carbon source. The best yield obtained was 2 g/L L-tyrosine from 10 g/L EG, achieving 50% of the theoretical yield (0.44 g L-tyrosine per g EG [**Fig. 1**]). The best L-tyrosine titer was 3 g/L (50 mL culture tube scale; 20 g/L EG). We also thoroughly compared EG and glucose as carbon substrate for L-tyrosine production and found that EG is a better substrate than glucose. A transcriptome analysis was done for understanding why EG was better than glucose for L-tyrosine production. It was further demonstrated that the strain can be modified to transform EG into a second aromatic amino acid, L-phenylalanine (1.5 g/L from 10 g/L EG). The engineered pathway for L-tyrosine production was also extended to produce *p*-coumaric acid (a valuable specialty chemical). The best strain produced 1 g/L *p*-coumaric acid from 10 g/L EG without any accumulation of L-tyrosine. Furthermore, we degraded waste PET bottles using acid hydrolysis and employed the resulting monomer EG as a substrate for L-tyrosine production, which yielded a similar L-tyrosine titer to that produced using commercial EG. The strains developed in this study may lead to more cost-effective processes for producing valuable aromatics including precursors for plastics, thereby setting the stage for establishing a circular bioeconomy in this area.

## 2. Results and Discussion

### 2.1. Developing a strain that can transform ethylene glycol into L-tyrosine

We recently found that simultaneous overexpression of propanediol oxidoreductase (encoded by *fucO*) and an aldehyde dehydrogenase (encoded by *aldA*) under a constitutive promoter substantially improved EG utilization in *E. coli* (Panda et al., 2021). The plasmid expressing *fucO* and *aldA* was termed as pEG03. A parallel study of our team (Pandit et al., 2021) found that two mutations to the N-terminus of FucO (I7L,L8V, encoded by *fucO\**) improved the enzyme’s oxygen tolerance. In this study, we first tested if combining the two strategies could further improve *E. coli’s* growth on EG. A new plasmid (pEG03*) was constructed by replacing *fucO* in pEG03 with fucO*. *E. coli* **MG1655DE3** harboring pEG03* grew slightly faster than that harboring pEG03 (**Supplementary Fig. 1**).

We next constructed *E. coli* **EGT01** by introducing pEG03* into *E. coli* **TPP15**, a strain we previously constructed for overproducing L-tyrosine from glucose (Ma et al., 2020). **TPP15** was derived from *E. coli* **MG1655DE3** by 1) deleting two genes from its genome (*ΔtyrR* for alleviating the transcriptional repression on L-tyrosine production; *ΔpheA* for eliminating formation of phenylalanine, a by-product; **MG1655DE3** *ΔtyrR ΔpheA* is named as **MG1655DE3TYR1**) and 2) introducing a plasmid (pTYR01) overexpressing feedback-resistant *aroG* and *tyrA* using a lac promoter.

In a chemically defined medium containing 10 g/L EG (Medium 1), optical density at 600 nm (OD_600_) of **EGT01** increased from 0.1 to 4.2 in 96 h and the strain produced 0.71 g/L L-tyrosine (**Fig. 2a)**. Medium 1 contained 1.6 g/L Complete Supplement Mixture (CSM, a mixture of amino acids and nucleobases) to facilitate the EG utilization. When we removed EG from Medium 1, OD_600_ of **EGT01** reached only 1 after 96 h and did not produce any L-tyrosine (**Fig. 2a**), suggesting that the produced L-tyrosine should be derived from EG. In Medium 1, **TPP15** could not utilize EG and did not produce L-tyrosine.

**Fig. 2.**
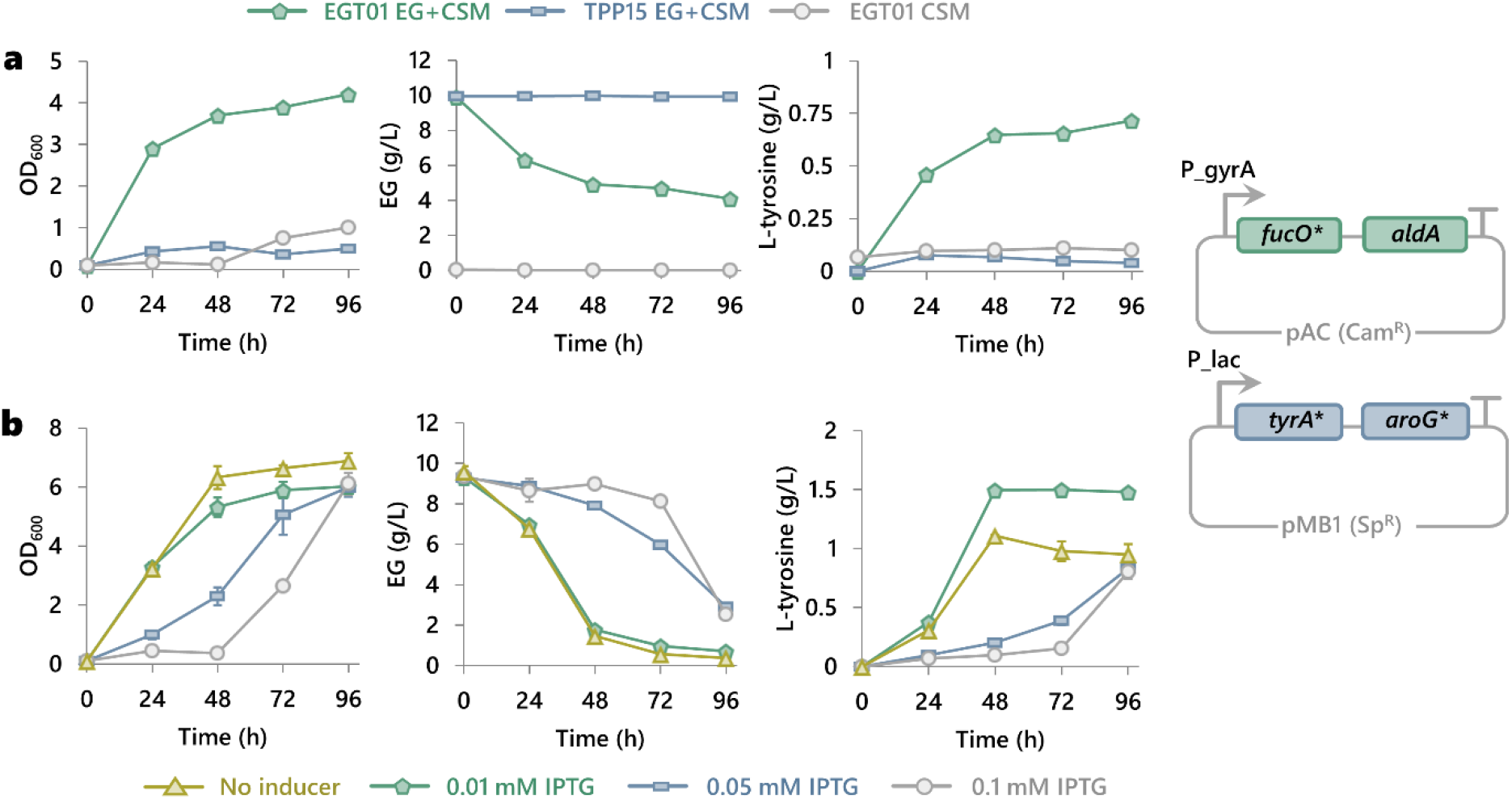
Production of L-tyrosine from ethylene glycol (EG) using engineered *E. coli*. (**a**) **EGT01** was grown in a chemically defined medium containing 10 g/L EG and 1.6 g/L complete supplement mixture (CSM). **EGT01** harboured two plasmids as shown. P_gyrA is a constitutive promoter. P_lac can be induced by using isopropyl β-D-1-thiogalactopyranoside (IPTG). 0.1 mM IPTG was added in early exponential growth phase. **TPP15** was grown in the same medium as a control. **TPP15** was the same to **EGT01** except **TPP15** does not have the EG-utilizing plasmid. There was also a control in which **EGT01** was grown in a medium without EG. (**b**) Effect of inducer concentration on L-tyrosine production. **EGT01** was grown in the same medium. Various IPTG concentrations (0-0.1 mM) were tested. IPTG was added upon inoculation. Due to leaky expression of the genes, 1 g/L L-tyrosine was produced even when no IPTG was added. 1.6 g/L CSM supplementation contained 0.1 g/L L-tyrosine, which was subtracted from the final titer in all the samples. Genotype of **EGT01** and other strains used in this study can be found in **Table 1**. The growth temperature was 30 °C. 10 mL culture was grown in 50 mL tube. Error bar indicates standard error (n=3).

In the above experiment, 0.1 mM isopropyl β-D-1-thiogalactopyranoside (IPTG) was added in early exponential growth phase to induce overexpression of *aroG* and *tyrA*. To avoid monitoring cell growth for adding the inducer, we tested adding the IPTG upon cell inoculation. The cells experienced a long lag phase possibly due to the burden caused by high expression level of the two genes (Dvorak et al., 2015). We reduced the expression level by using lower concentrations of IPTG (0.05 mM and 0.01 mM; added upon cell inoculation). Cell growth was gradually improved with the decrease of the IPTG concentration. More importantly, the L-tyrosine titer and productivity were also substantially improved. The culture with 0.01 mM IPTG produced 1.5 g/L L-tyrosine in 48 h (**Fig. 2b**). We further decreased the expression level of the two genes by using the leaky expression (without adding IPTG). A lower L-tyrosine titer was achieved. We replaced the promoters for driving the EG utilization operon and/or the L-tyrosine production operon with auto-inducible or constitutive promoters (pJ23114, P_thrC3, P_gyrA), however, no substantial improvement in L-tyrosine titer was observed (**Supplementary Figs. 2-3**). We continued to use **EGT01** in the subsequent experiments with 0.01 mM IPTG induction upon inoculation.

### 2.2. Improving L-tyrosine production through medium optimization and process engineering

We next increased the EG concentration in Medium 1 from 10 g/L to 20 g/L (Medium 2). In Medium 2, the cells stopped to grow at 48 h, at which 1.5 g/L L-tyrosine was produced and ~10 g/L EG was left (**Fig. 3a**). In the next 48 h, ~6 g/L of EG was assimilated with no L-tyrosine production. Through the fermentation, no organic acids (e.g., acetate, glycolate) and other soluble, frequently encountered by-products were detected (detection limit: 0.1 g/L). It is possible that a large fraction of the consumed EG was fully oxidized into carbon dioxide (CO_2_) when it is not transformed into L-tyrosine or biomass, especially during the stationary phase (Panda et al., 2021). It was suggested that L-tyrosine production was growth-associated (Lutke-Eversloh and Stephanopoulos, 2007). The hypothesis was further supported by two experiments described in **Supplementary Note S1** and **Supplementary Fig. 4**.

**Fig. 3.**
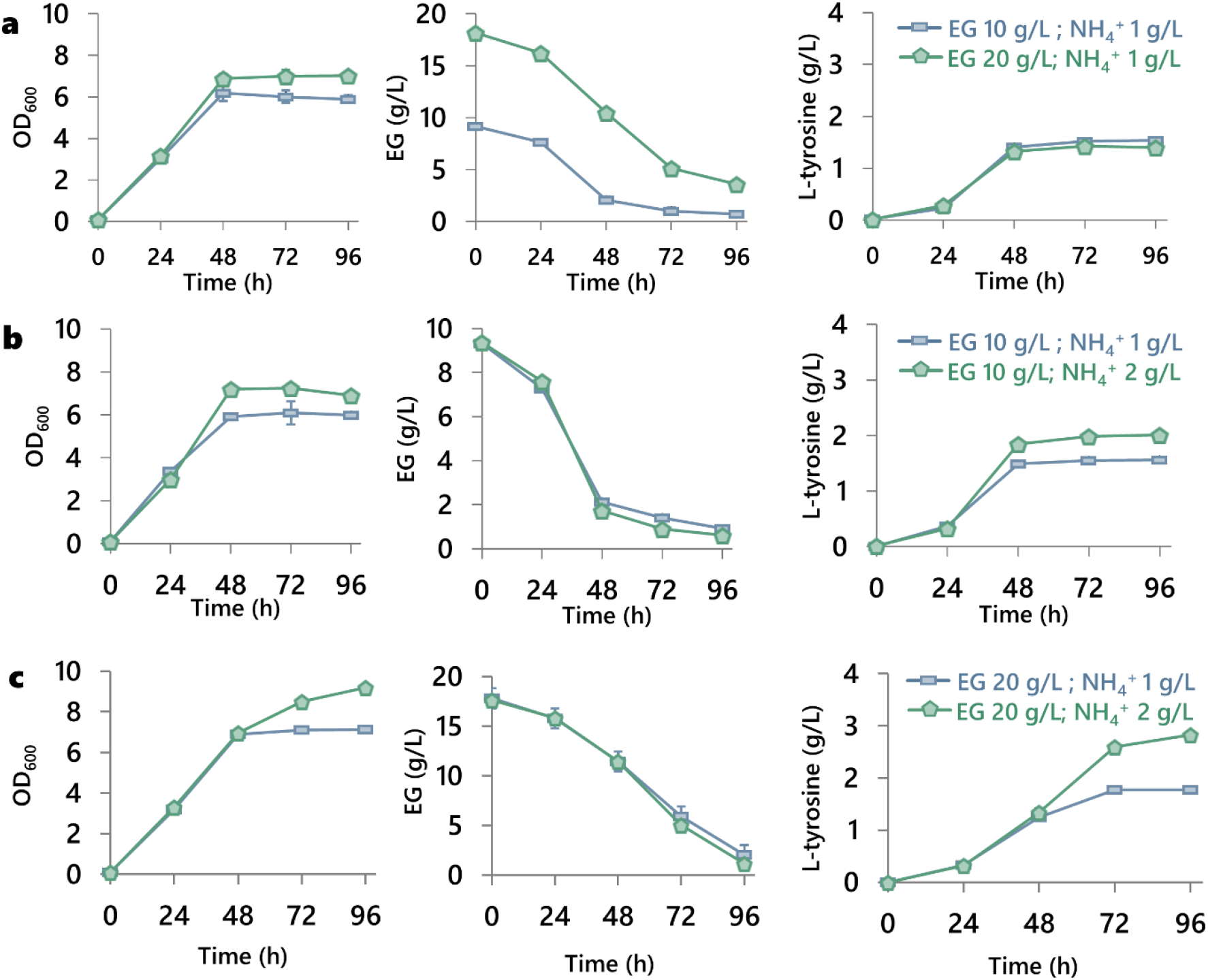
Improving L-tyrosine production using medium optimization. (**a**) At low ammonium concentration (1 g/L), increasing EG concentration from 10 g/L to 20 g/L did not improve L-tyrosine production. (**b**) At low EG concentration (10 g/L), increasing ammonium concentration from 1 g/L to 2 g/L did not improve L-tyrosine production. (**c**). At high EG concentration (20 g/L), increasing ammonium concentration from 1 g/L to 2 g/L substantially improved L-tyrosine production. *E. coli* **EGT01** was induced with 0.01 mM IPTG upon inoculation. The temperature was 30 °C. 10 mL culture was grown in 50 mL tube. Error bar indicates standard error (n=3).

We further hypothesized that the nitrogen and/or phosphorous sources were depleted by 48 h during cell growth in Medium 2, when the initial EG concentration was increased from 10 g/L to 20 g/L. Therefore, we systematically evaluated the effect of ammonium and phosphate concentrations on L-tyrosine production. When the ammonium concentration of Medium 1 (EG concentration: 10 g/L) was increased from 1 g/L to 2 g/L, the L-tyrosine titer was increased from 1.5 g/L to 2 g/L. (**Fig. 3b**). The final cell density was also slightly increased. When the ammonium concentration in Medium 2 (EG concentration: 20 g/L) was increased from 1 g/L to 2 g/L, there was substantial improvement in the final cell density and L-tyrosine production. 3 g/L L-tyrosine was produced from 17 g/L EG (**Fig. 3c**).

### 2.3. Comparing L-tyrosine production from EG and glucose

Glucose is one of the most frequently used carbon sources for *E. coli*. We compared glucose and EG for L-tyrosine production. **EGT01** was grown on 10 g/L EG or glucose. **EGT01** could not grow on EG without CSM, so all the EG mediums contained 1.6 g/L CSM. Half of the glucose mediums contained CSM (1.6 g/L) to investigate its effect on L-tyrosine production. All the cultures contained 2 g/L NH4^+^.

We used two induction conditions: 1) the best induction condition when EG was used as carbon source (0.01 mM IPTG added upon inoculation); 2) the commonly used induction condition for glucose (0.1 mM IPTG added during the early exponential phase) (Ma et al., 2020). The best L-tyrosine titer obtained from the glucose media was only 1.2 g/L (96 h, **Fig. 4**), corresponding to ~23% of the theoretical yield (0.55 g L-tyrosine per g glucose) (Juminaga et al., 2012), which was much lower than that with EG as a substrate (2 g/L). Furthermore, the cells accumulated a large amount of acetate in the glucose mediums whereas no byproducts were observed in the EG culture. The results suggested that EG is a better carbon source than glucose for L-tyrosine production.

**Fig. 4.**
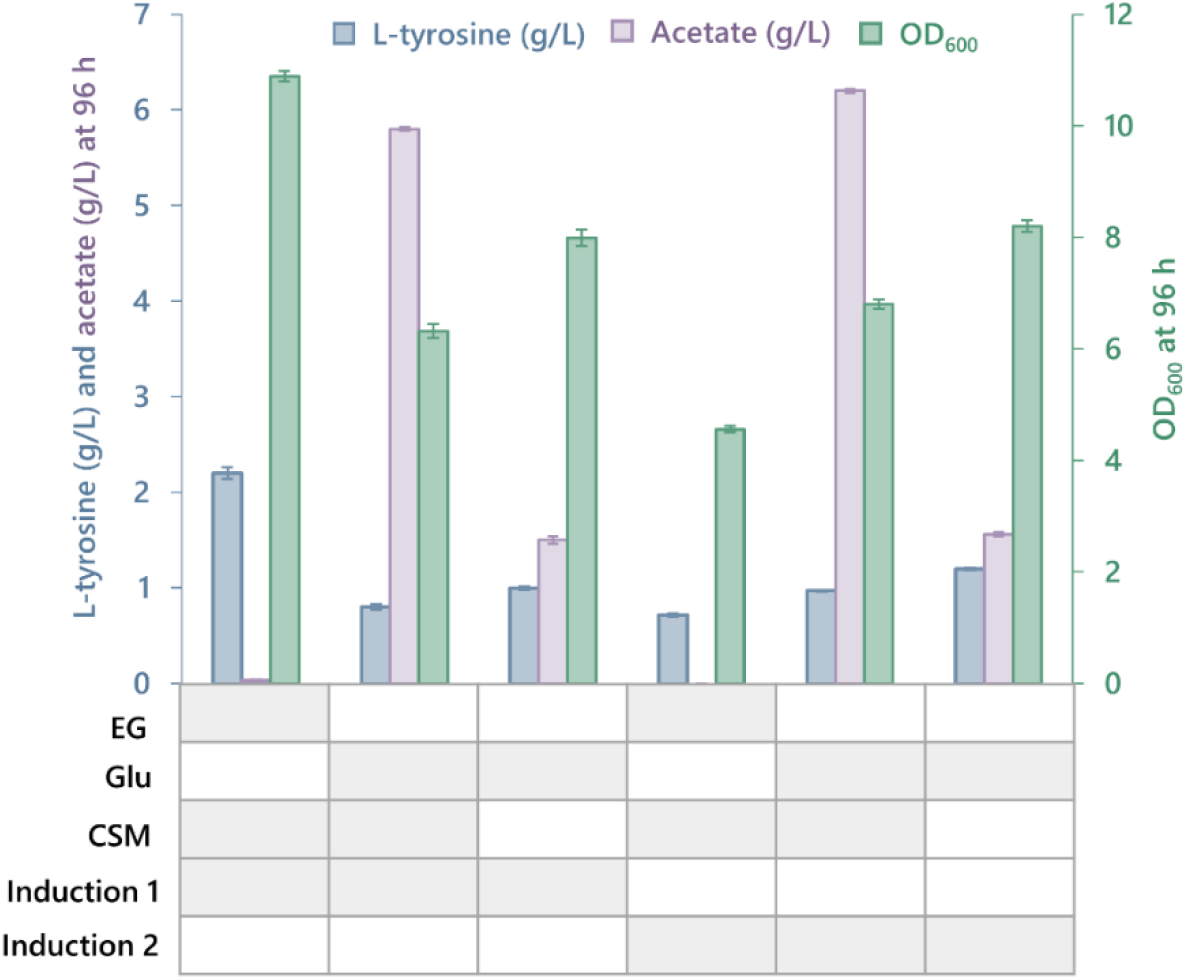
Comparing EG and glucose as substrate for producing L-tyrosine. **EGT01** was grown in chemically defined medium containing 10 g/L EG or 10 g/L glucose. Some cultures contained 1.6 g/L CSM. Two induction conditions were compared: 0.01 mM IPTG added upon inoculation; 0.1 mM IPTG during early exponential phase (OD_600_ = ~0.5). The culture volume was 10 mL. The culture vessel was 125 mL shake flask. The growth temperature was 30 °C. Error bar indicates standard error (n=3).

### 2.4. Comparative transcriptome analysis of L-tyrosine-producing strain growing on EG or glucose

To understand the metabolism of **EGT01** when it grew on EG and why EG is a better substrate than glucose for L-tyrosine production, we extracted total RNA during its early exponential growth phase on 10 g/L EG or 10 g/L glucose. The total RNA was analyzed using Illumina sequencing after a standard sample preparation procedure. *fucO* and *aldA* were upregulated by approximately 2-fold when glucose in the medium was replaced with EG (**Fig. 5**). These two genes were expressed from both plasmids and genome to transform EG into glycolate. Further data analysis revealed that most of the transcripts were contributed by the plasmid, suggesting the promoter used to express *fucO* and *aldA* on the plasmid (P_gyrA) may be repressed by glucose or activated during growth on EG. Glycolate was transformed into glyoxylate by glycolate dehydrogenase, whose encoding genes (*glcDEF*) were upregulated by >100-fold during cell growth on EG. The genes in the glycerate pathway (*gcl, glxR and glxK/garK*; encoded in the genome) were also upregulated by >100-fold. These genes’ products collectively transform glyoxylate into 2-phosphoglycerate (2PG), which provides an entry point into glycolysis/gluconeogenesis. These large gene expression upregulations were expected because the glycerate pathway was the only way of assimilating glyoxylate into the central metabolism under the condition (Panda et al., 2021).

**Fig. 5.**
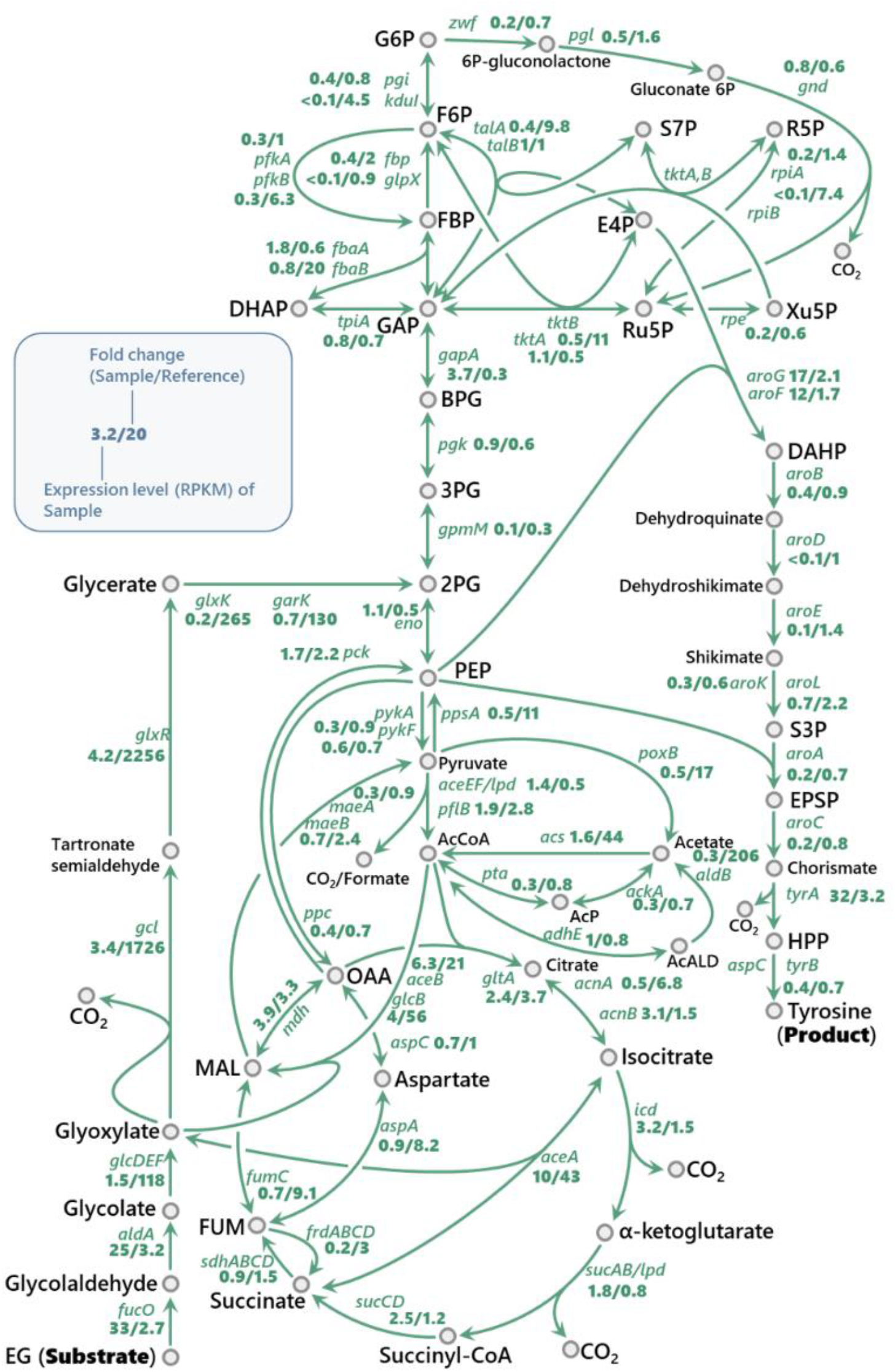
Comparative transcriptome analysis of **EGT01** when it grew on EG or glucose. Key metaboates in central metabolism are denoted by using commonly used abbreviations. **EGT01** was cultured in a chemically defined medium containing either 10 g/L EG or 10 g/L glucose. Total RNA was extracted at the early exponential growth phase (OD_600_ = ~0.5). The cells were induced with 0.01 mM IPTG upon inoculation. The temperature was 30 °C in all the experiments. 10 mL culture was grown in 125 mL shake flask. Error bar indicates standard error (n=3). The blue box is the legend to explain the data presentation style. RPKM: Reads Per Kilobase of transcript, per Million mapped reads.

Although glyoxylate could also enter the TCA cycle through malate synthase (GlcB or AceB) or isocitrate lyase (AceA), these reactions would need to borrow acetyl-CoA or succinate, respectively. When acetyl-CoA and succinate are returned through oxidative decarboxylation reactions, the assimilated carbon atoms would be completely lost as CO_2_ (Panda et al., 2021). These two pathways could provide the reducing equivalents and energy to support cell growth and product formation but cannot contribute carbon atoms. *aceA*, *aceB* and *glcB* were upregulated by at least 20-fold, suggesting that both pathways may be operating during the cell growth on EG.

2PG can be dehydrated by enolase (Eno) to form PEP, a critical building block of L-tyrosine. Through the glycolysis PEP could become pyruvate, which is primarily transformed into acetyl-CoA by pyruvate dehydrogenase (PDH) complex during cell growth on glucose. When cells growing on EG, we observed activation of two alternative routes to transform pyruvate into acetyl-CoA: *poxB* and *acs* were upregulated by 17-fold and 44-fold respectively (PoxB oxidatively decarboxylates pyruvate into acetate and Acs activates acetate into acetyl-CoA); pflB was upregulated by approximately 2-fold (PflB splits pyruvate into acetyl-CoA and formate in one step). The activation of the two pathways suggests that the PDH complex may have been partially or completely inactivated during the growth on EG (the transcription of PDH genes was halved during the growth on EG and some PDH proteins may be inhibited by NADH or damaged by reactive oxygen species [ROS]). This would facilitate accumulation of PEP, resulting in a higher L-tyrosine titer. Furthermore, we noticed that *ppsA* (PEP synthase) was upregulated by 11-fold, indicating increased PEP availability for L-tyrosine production in EG medium.

Through the above analysis, we at least could suspect that the improved L-tyrosine production by using EG should be partly due to 1) the elevated expression of key enzymes (aroG and tyrA), and 2) rewiring of the central metabolism for pushing flux towards the shikimate pathway.

### 2.5. Converting EG into other aromatic chemicals

To prove the concept that EG can be converted into different aromatic chemicals, we engineered *E. coli* to synthesize L-phenylalanine from EG. L-phenylalanine is the key ingredient in the synthesis of many edible products, such as artificial sweetener aspartame (Rodriguez et al., 2014). Chorismate is a common precursor of L-phenylalanine and L-tyrosine. It can be rearranged and decarboxylated to form phenylpyruvate, with PheA as the catalyst. Phenylpyruvate can be converted into L-phenylalanine through transamination by using L-glutamate as the amino group donor (**Fig. 6a**).

**Fig. 6.**
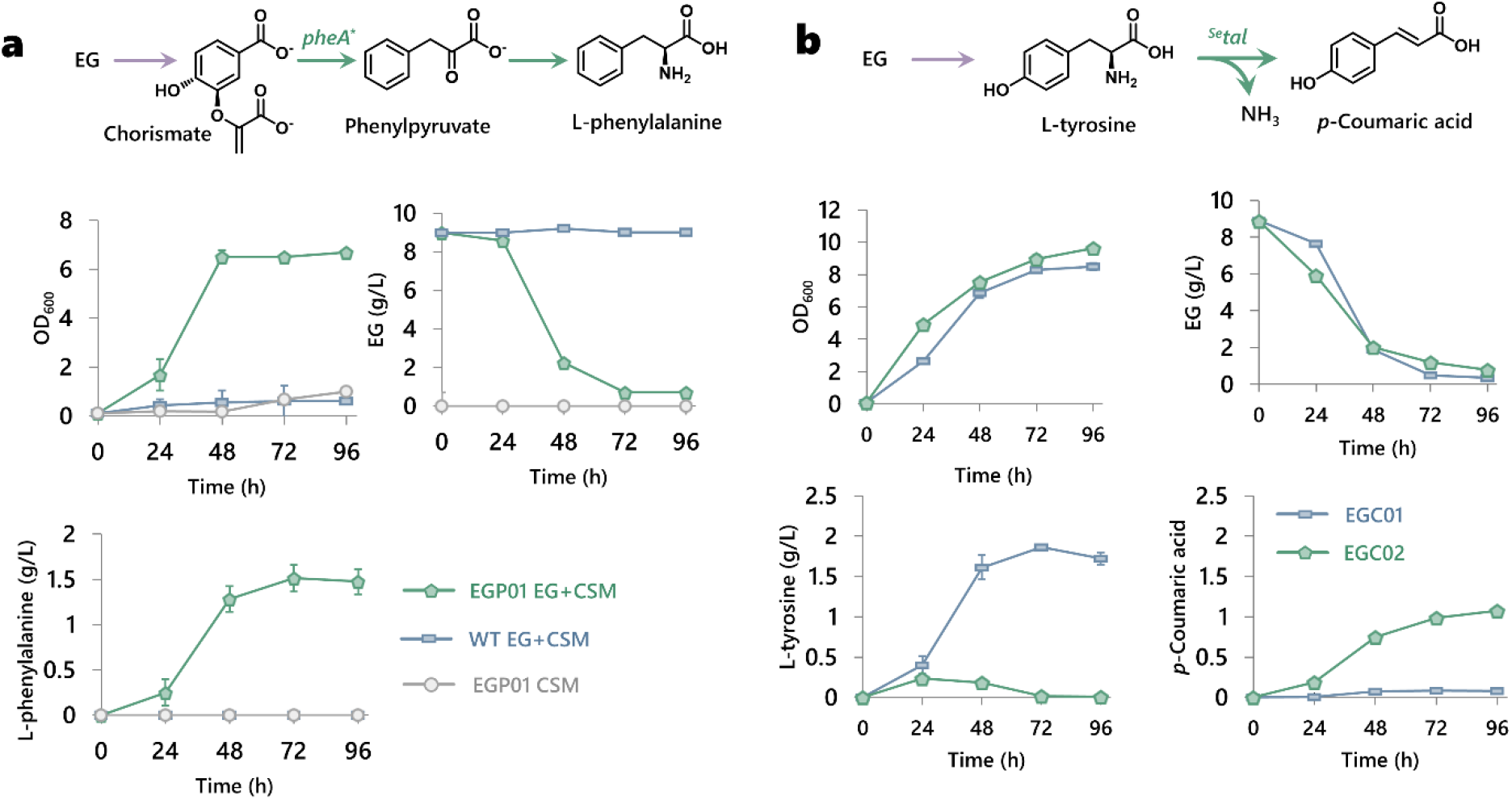
Production of L-phenylalanine and p-coumaric acid from ethylene glycol (EG) using engineered *E. coli*. (**a**) **EGP01** was a strain engineered to produce L-phenylalanine from EG. It had a plasmid to utilize EG and a plasmid to produce L-phenylalanine. **EGP01** was grown in a chemically defined medium containing 10 g/L EG and 1.6 g/L complete supplement mixture (CSM). **WT** is a control strain which was the same to **EGP01** except that it did not have the EG-utilizing plasmid. **EGP01** was also grown in an EG-free medium as a control. 0.01 mM IPTG was added upon cell inoculation. (**b**) **EGC01** was first engineered to produce p-coumaric acid (pCA) from EG. The critical gene ^Se^*tal* was inserted after the third gene into the L-tyrosine operon, far from the promoter. After the pCA titer was found to be low, we constructed **EGC02**, in which ^Se^*tal* was expressed in a single-gene operon. The growth temperature was 30°C. Error bar indicates standard error (n=3).

The feedback resistant mutant of PheA* (Thr326Pro) and AroG* were overexpressed under a P_lac promoter. The resulting plasmid (pPHE01) and plasmid pEG03* were introduced into a **MG1655DE3** derivative containing the following two deletions: *ΔtyrR* for alleviating the transcriptional repression on phenylalanine production and *ΔtyrA* for minimizing the carbon loss via the L-tyrosine-producing pathway. The resulting strain (**EGP01**) produced 1.5 g/L L-phenylalanine from 10 g/L EG and 1.6 g/L CSM (**Fig. 6a**). The control strain (without pEG03* plasmid) did not produce any L-phenylalanine in the same medium. When grown solely on CSM without any EG, **EGP01** did not produce any L-phenylalanine. 1.6 g/L CSM supplementation contained 0.1 g/L L-phenylalanine, which was subtracted from the final titer in all samples. These results supported that the produced phenylalanine was derived from EG.

*p*-Coumaric acid (pCA) is a valuable specialty chemical and can be derived into many secondary metabolites, including flavonoids (Li et al., 2018). We therefore also explored if it is possible to efficiently synthesize pCA from EG. We engineered **EGT01** to express tyrosine ammonia lyase (Tal, encoded by *^Se^tal*) which could transform L-tyrosine into pCA (the new strain: **EGC01**, **Fig. 6b**). The gene encoding Tal was inserted after *tyrA\** and *aroG\** in one operon. In Medium 1 (10 g/L EG), **EGC01** produced 0.07 g/L p-coumarate and 1.7 g/L L-tyrosine. It was clear that TAL catalyzed the rate-limiting step. The low TAL activity could be due to the fact that *tal* was the third gene in the operon, far from the promoter. We rearranged the two plasmids in **EGC01** to overcome this challenge.

We first created a plasmid (pEGTYR01) carrying both the EG utilization operon (P_gyrA-*fucO\**-*aldA*) and the L-tyrosine production operon (P_lac-*tyrA\**-*aroG\**), and confirmed that introducing pEGTYR01 into **MG1655DE3TYR1** could enable the new strain (**EGT08**) to produce 2 g/L L-tyrosine in Medium 1. We then constructed another plasmid (pCA02) only expressing Tal under the lac promoter. When pCA02 were introduced into **EGT08**, the new strain (**EGC02**) produced ~1 g/L p-coumarate with no L-tyrosine accumulation (**Fig. 6b**).

Although the pCA titer was among the highest reported titers in the literature (shake flask scale, from 10 g/L EG), the titer was approximately 50% of the L-tyrosine titer obtained by using **EGT08** (the parent strain) in the same medium. We hypothesized that introducing pCA02 into **EGT08** may have compromised its ability of producing L-tyrosine. To test this hypothesis, we cultured **EGT08** in Medium 1 for 96 h, removed **EGT08** by centrifugation, and used the spent medium (containing 2 g/L L-tyrosine) to resuspend a Tal-expressing strain (**CA01**) at various cell concentrations (OD_600_: 0.6 to 6). The **CA01** cells had been cultured separately in LB medium for Tal expression. The resuspended cells were incubated at 30 °C / 250 rpm for 96 h, which was defined as the second stage. Under the best condition, 1 g/L of *p*-coumarate was obtained in the second stage in which all the L-tyrosine was consumed (**Supplementary Fig. 5**). The results suggest that L-tyrosine and/or pCA may not be chemically stable in the culture of **CA01**, and that **EGC02** may have the same ability of producing L-tyrosine as **EGT08**.

### 2.6. Production of L-tyrosine from PET wastes

To demonstrate the production of L-tyrosine from PET waste, we used post-consumer PET bottles. Their neck portions, which have been reported to have less crystallinity than other sections of the bottles (Lu et al., 2022), were cut, and depolymerized using acid hydrolysis (Mancini and Zanin, 2007). The EG was separated from terephthalic acid (TPA) and residual PET, and subsequently desalted. We obtained 0.65 g of EG from 5.82 g of PET waste, achieving 40% of the theoretical recovery yield (maximum theoretical yield is 0.27 g EG per g PET) (**Fig. 7a**). The obtained EG was used to replace commercial EG in the chemically defined medium reported in **Section 2.4**. When the medium was seeded with **EGT01**, the cells grew normally as the commercial EG control, and produced similar quantity of L-tyrosine as the control (**Fig. 7b**). The results proved the concept that plastic waste can be transformed into food ingredients using chemical hydrolysis and the *E. coli* strain we engineered in this study.

**Fig. 7.**
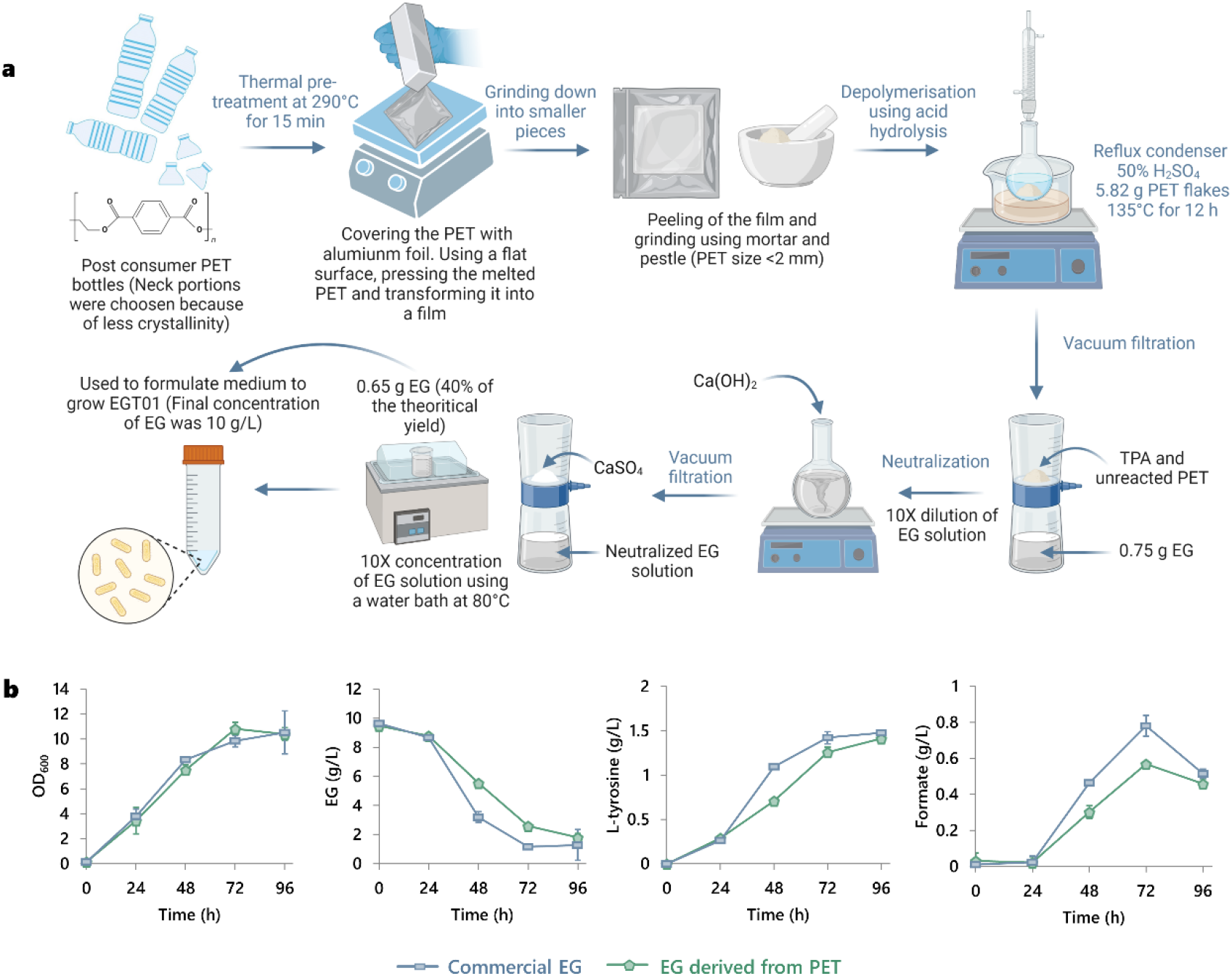
Production of L-tyrosine from PET degradation. (**a**) Post-consumer PET bottles were depolymerized using acid hydrolysis to generate their monomers (EG and TPA). The liquid containing EG was filtered and desalted to formulate the medium for growing **EGT01**. PET: Polyethylene terephthalate. TPA: Terephthalic acid. (**b**) **EGT01** was grown in a medium containing 10 g/L PET-derived EG. A medium formulated using commercial EG at an equivalent concentration was used as a control. 2 g/L NH_4_Cl was used. **EGT01** was induced with 0.01 mM IPTG upon inoculation. 10 mL of culture was grown in 50 mL tube. The growth temperature was 30 °C. Error bar indicates standard error (n=3).

We chose to use the conventional acid PET hydrolysis in this study because it did not require special reagent/equipment and could achieve decent hydrolysis efficiency. A major limitation is that the use of high concentration of sulfuric acid (50 wt%) required tedious and wasteful purification steps to desalt the product. Without desalting, *E. coli* would not be able to grow due to the high salt concentration. The hydrolysis step could be improved in a few possible ways in the future. PETase could complete the hydrolysis under neutral conditions (Lu et al., 2022) and improving the related processes is being intensively investigated by many companies and academic labs. Some abiotic catalytic processes are also promising. For example, close to theoretical recovery yield was achieved when TPA was used as a catalyst under 220 °C (Yang et al., 2021). Since TPA is also a product and can be easily separated from EG, the method could yield suitable EG stream for downstream fermentation processes. The obtained TPA could be used to hydrolyze more PET waste and/or used as substrate to synthesize new PET products.

## 3. Conclusions

This study demonstrated that *E. coli* can be engineered to produce L-tyrosine from PET-derived EG, a sustainable feedstock. A wide range of other valuable aromatic chemicals (L-phenylalanine and *p*-coumaric acid) could also be produced from EG. Hence, our study sets the stage for expanding the range of products that can be made from EG, given that EG can be transformed into intermediates in the central metabolism within two steps. EG was found to be a better substrate than glucose to produce L-tyrosine. Future studies may improve the production of the aromatics through manipulation of the central metabolism and/or introducing new pathways to assimilate EG. In addition, more carbon-efficient EG utilization pathways can also be evaluated to further enhance yields of the products from EG. These efforts would collectively contribute to developing a more circular bioeconomy, where waste streams can be upcycled into valuable chemicals using EG as an intermediate.

## 4. Materials and methods

### 4.1. Plasmids and strains

All the chemicals used in this study were purchased from Sigma-Aldrich unless otherwise specified. D-glucose, L-tyrosine and M9 medium broth powder were purchased from Bio Basic Asia Pacific Pte Ltd. Complete Supplement Mixture (CSM) powder was purchased from Sunrise Science Products. The strains and plasmids used in this study are provided in **Table 1**. All the plasmids were constructed according to the GT standard (Ma et al., 2019). The Addgene IDs of the key plasmids are provided in **Table 1**. The sequences and annotations can be retrieved using the Addgene IDs. The primers used in this study were synthesized by Integrated DNA Technologies (IDT) (their sequences are included in **Supplementary Table 1**). The C1000 Thermal cycler (Bio-Rad) was used to perform PCR. Q5^®^ High-Fidelity 2×Master Mix (New England Biolabs) was used in all the PCR reactions. The sequences of all the constructed plasmids were verified using Sanger sequencing (Service provider: Bio Basic Asia Pacific Pte Ltd, Singapore).

**Table 1.** Strains constructed in this study.

| Strain | Parent strain | Plasmid in the strain/modifications |
| --- | --- | --- |
| EG05 | MG1655DE3 | pEG03* (P<sub>gyrA</sub> - *fucO* *- *aldA* -p15A-cam, [185811]) |
| TPP15 | MG1655DE3TYR1 | pTYR01 (P<sub>lac</sub> - *tyrA* *- *aroG* *-pMB1-spec, [185812]) |
| EGT01 | MG1655DE3TYR1 | pEG03*<br>pTYR01 |
| EGT02 | MG1655DE3TYR1 | pEG03*<br>P<sub>gyrA</sub> - *tyrA* *- *aroG* *-pMB1-spec |
| EGT03 | MG1655DE3TYR1 | pEG03*<br>P<sub>thrC3</sub> - *tyrA* *- *aroG* *-pMB1-spec |
| EGT04 | MG1655DE3TYR1 | pEG03*<br>P<sub>J23114</sub> - *tyrA* *- *aroG* *-pMB1-spec |
| EGT05 | MG1655DE3TYR1 | P<sub>proX</sub> - *fucO* *- *aldA* -p15A-cam<br>pTYR01 |
| EGT06 | MG1655DE3TYR1 | P<sub>J23114</sub> - *fucO* *- *aldA* -p15A-cam<br>pTYR01 |
| EGT07 | MG1655DE3TYR1 | P<sub>J23114</sub> - *fucO* *- *aldA* -p15A-cam<br>P<sub>J23114</sub> - *tyrA* *- *aroG* *-pMB1-spec |
| EGT08 | MG1655DE3TYR1 | pEGTYR01 (P<sub>gyrA</sub> - *fucO* *- *aldA* -P<sub>lac</sub> - *tyrA* *- *aroG* *-p15A-cam) |
| EGP01 | MG1655DE3PHE1 | pEG03*<br>pPHE01 (P<sub>lac</sub>-pheA*-aroG*-pMB1-spec) |
| EGC01 | MG1655DE3TYR1 | pEG03*<br>pCA01 (P<sub>lac</sub>-tyrA*-aroG*-tal-pMB1-spec) |
| EGC02 | MG1655DE3TYR1 | pEGTYR01<br>pCA02 (P<sub>lac</sub>-tal-pMB1-spec) |
| CA01 | MG1655DE3TYR1 | pCA02 |

pMB1 and p15A are the high copy number and medium copy number replication origins respectively; spec and cam are spectinomycin and chloramphenicol resistance cassettes respectively. **MG1655DE3TYR1**: **MG1655DE3** ΔpheA ΔtyrR. **MG1655DE3PHE1**: **MG1655DE3** ΔtyrA ΔtyrR. The texts in brackets are key features of the associated plasmid. 185811 and 185812 are Addgene IDs.

To construct pEG03* plasmid, a *fucO* mutant (Ile7Leu, leu8val) and *aldA* were overexpressed under constitutive promoter P_gyrA. pTYR01 and pEG03* were introduced into *E. coli* **MG1655DE3TYR1**, by using the standard heat-shock method, creating the strain **EGT01**.

To construct **EGT02**, **EGT03** and **EGT04**, the P_lac promoter of pTYR01 was replaced with P_gyrA, P_thrC3 (auto inducible promoter (Anilionyte et al., 2018)) and P_J23114 (Weak constitutive promoter [registry of standard biological parts collections, Anderson promoters]) respectively. To construct **EGT05** and **EGT06**, P_gyrA in pEG03* was replaced with P_ProX (strong constitutive promoter) and P_J23114 respectively. To construct **EGT07**, P_gyrA and P_lac promoter in both pEG03* and pTYR01 were replaced with P_J23114. When EG utilization operon (P_gyrA-fucO*-aldA) and L-tyrosine production operon (P_lac-tyrA*-aroG*) were combined into a single plasmid, the resulting plasmid was named as pEGTYR01. pEGTYR01 was introduced into **MG1655DE3TYR1** to create **EGT08**.

The L-phenylalanine producing strains were derived from **MG1655DE3** by inactivating *tyrR* and tyrA (**MG1655DE3PHE1**). To construct pPHE01 plasmid, *pheA\**(T326P) and *aroG\**(D146N) were over-expressed under P_lac promoter. pPHE01 was introduced into **MG1655DE3PHE1**, resulting in the new strain, **EGP01**.

To construct pCA01, *tal* from from *Saccharothrix espanaensis* was cloned as the third gene in the operon after aroG* and tyrA* under a P_lac promoter. pEG03* and pCA01 were introduced into **MG1655DE3TYR1** to create the strain **EGC01**. To create pCA02, only ^Se^*tal* was overexpressed in a single gene operon with a high copy number replication origin (pMB1) under P_lac promoter. pCA02 was introduced into **EGT08**, creating **EGC02**. To create **CA01**, pCA02 was introduced into **MG1655DE3TYR1.**

### 4.2. Growth media

LB medium was used to prepare seed culture. We used modified M9 media containing X g/L ethylene glycol (X was varied between 10 and 20 as specified in the body text), Y g/L ammonium chloride (NH_4_Cl, Y was varied between 1 and 2), 6.78 g/L disodium phosphate (Na_2_HPO_4_), 3 g/L monopotassium phosphate (KH_2_PO_4_), 0.5 g/L sodium chloride (NaCl), 0.17% (V/V) K3 master mix, and 1.6 g/L CSM. The K3 master mix was prepared by mixing 2.5 mL of 0.1 M ferric citrate solution (autoclaved), 1 mL of 4.5 g/L thiamine solution, 3 mL of 4 mM of Na_2_MoO_4_ (autoclaved), 1 mL of 1 M MgSO_4_ solution (autoclaved) and 1 mL of 1000X K3 trace elements stock solution. The 1000X K3 trace elements stock solution contained 5 g/L CaCl_2_·2H_2_O, 1.6 g/L MnCl_2_·4H_2_O, 0.38 g/L CuCl_2_·2H_2_O, 0.5 g/L CoCl_2_·6H_2_O, 0.94 g/L ZnCl_2_, 0.03 g/L H_3_BO_3_, 0.4 g/L Na_2_EDTA·2H_2_O. The pH of the medium was adjusted to 7. Antibiotics (25 μg/L chloramphenicol or 50 μg/L spectinomycin) was used to create selection pressure to maintain the plasmids. The antibiotic resistance of each strain is described in **Table 1**. IPTG (0.01-0.1 mM) was added to induce the protein expression, when inducible promoter (P_lac) was used.

### 4.3. Culture conditions

Single colony was used to inoculate LB that contains the antibiotics. The seed culture was incubated at 37 °C/250 rpm overnight. The overnight culture was centrifuged at 4,000 g for 10 mins and washed two times with ultrapure water. The cell pellet was used to inoculate the modified M9 medium in 50 mL tubes or 125 mL shake flasks (as specified in the figure captions). The initial optical density at 600 nm (OD_600_) was 0.1. The antibiotic was added to maintain the L-tyrosine producing plasmids, while no antibiotic was added for EG plasmids since any cell losing the plasmid could not grow on EG. The culture was incubated at 30 °C/250 rpm for 72 hours. The cell culture was induced by IPTG (0.01-0.1 mM) either at the inoculation or at the early exponential phase as indicated in the relevant sections, when P_lac promoter was used.

When the sequential culture strategy was used for pCA production, **EGT08** was cultured until 96 h to produce 2 g/L L-tyrosine. At 96 h, cells were centrifuged at 4,000 g for 10 mins and washed. **CA01** had been cultured separately in LB medium for Tal expression (induction: 0.01 mM IPTG upon inoculation). The spent medium containing 2 g/L L-tyrosine obtained in the first stage was used to resuspend **CA01** at different concentrations (OD_600_ 0.6-6). The resuspended cells were incubated for 96 h at 30 °C.

### 4.4. Quantification of cell density and metabolites

OD_600_ at indicated time points were measured using a microplate reader (Tecan infinite M200). The raw data were converted into standard OD_600_ units using a standard curve.

To dissolve the L-tyrosine crystals, 50 μL of 6 M HCL was added to 500 μL of cell culture and the culture was mixed by vortex (37 °C for 30 min). The mixture was then centrifuged at 13,000 g for 5 min. The supernatant was filtered and analyzed using a high-performance liquid chromatography (HPLC). The column was Agilent ZORBAX Eclipse plus C18 (3.5 μm, 4.6X100 mm). An isocratic flow was used with a flow rate of 0.7 mL/min. The mobile phase consisted of 89.9% (v/v) water, 10% (v/v) acetonitrile, and 0.1 % (v/v) trifluoracetic acid. The column temperature was 30 °C. The detector was a UV detector with the wavelength set at 275 nm for quantifying L-tyrosine.

To measure the *p*-coumaric acid, 50 μL of 6 M HCL was added to 500 μL of cell culture and the culture was mixed by vortex. 350 μL of acetonitrile was then mixed with 150 μL of the acidified culture. The mixture was centrifuged at 13,000 g for 5 min. The supernatant was filtered and analyzed using HPLC. The column was the same to what was described above. An isocratic flow was used with a flow rate of 0.8 mL/min. The mobile phase consists of 79.9% (v/v) water, 20% (v/v) acetonitrile, and 0.1 % (v/v) trifluoracetic acid. The column temperature was 30 °C. The detector was a UV detector with the wavelength set at 310 nm for quantifying *p*-coumaric acid.

To quantify EG, glycolate and other fermentation by-products (acetate, lactate, ethanol, glycerol, succinate, and formate), 0.2 mL of cell culture was collected at the indicated time points. The supernatant was centrifuged for 5 min at 12,000 g, and the supernatant was filtered using a nylon syringe filter with a pore size of 0.22 μm and a diameter of 13 mm (IT Technologies Pte Ltd). 5 μL of the obtained filtered supernatant was analyzed using HPLC. The column used was Aminex HPX-87H (300X7.8 mm, Bio-Rad). 5 mM H2SO4 was used as mobile phase with a flow rate of 0.7 ml/min (isocratic flow). The compounds were detected using a refractive index detector (RID). A calibration curve was obtained for each compound. Products were identified based on their retention time.

### 4.5. Comparative Transcriptome Analysis

**EGT01** was grown on EG or glucose with CSM supplementation, induced with 0.01 mM IPTG at the inoculation. Total RNA was extracted from the fresh culture at early exponential phase (OD_600_ 0.5-1), according to the manufacture’s protocol (Thermo Scientific GeneJET RNA purification kit, K0731). The RNA concentrations were measured using Nanodrop. The samples were sequenced using the Illumina technology after a standard sample prep procedure (PE150; Service Provider: Genewiz, China). The FASTQ data were processed using an in-house developed MATLAB App.

### 4.6. Maximum Theoretical Product Yield Calculations

The maximum theoretical yield of L-tyrosine from both EG and glucose as main carbon sources, respectively, were calculated using flux balance analysis (FBA). FBA was performed using the COBRApy toolbox on Python 3.7.9 using the GLPK solver. The core model of *E. coli* was modified to include EG assimilation and L-tyrosine biosynthetic metabolic steps (Orth et al., 2010). To calculate the maximum theoretical yield of L-tyrosine from EG or glucose, the respective carbon source was designated as the sole substrate and FBA was performed without constraints on by-product fluxes and oxygen uptake.

### 4.7. PET depolymerization by acid hydrolysis

A garden cutter was used to cut out the neck portions of post-consumer PET plastic bottles (reported to have less crystallinity compared to other parts of the bottles). The neck portions were then covered with aluminum foil and undergone thermal pretreatment for 30 mins on a hot plate at 290 °C (Lu et al., 2022). The molten PET was pressed into a film using a 6061-aluminum alloy block. After cooling, the film was peeled off, and pulverized into small (< 2 mm) pieces with a mortar and pestle.

The resulting PET flakes were depolymerized using acid hydrolysis. 50 mL of 50% sulphuric acid containing 5.82 g of PET flakes was agitated using a magnetic stirrer for 12 h. The reaction temperature was 135 °C and was achieved using a 500 mL round bottom flask with a reflux condenser (Mancini and Zanin, 2007). The solid component containing TPA and the unreacted PET were filtered out using a vacuum filter (Corning 431098 1L filter with 0.22 μm polyether sulfone membrane). The filtrate was diluted 10 times, and Ca(OH)2 powder was slowly added to the solution until pH > 7. In the above step, the solution was agitated continuously in a round bottom flask using magnetic stirrer. The neutralized solution was filtered using a vacuum filter (Corning 431098 1L filter with 0.22 μm polyether sulfone membrane) to remove the salts. The filtrate was concentrated 10 times at 80 °C, then cooled at ambient temperature before 0.22 μm filter sterilization. The resulting EG solution was used to formulate the medium for growing **EGT01** (final EG concentration in the culture medium was 10 g/L).

## Data availability

All the data supporting the findings of this study are included in this paper and its supplementary information file.

## Acknowledgement

This work was financially supported by research grants from Singapore National Research Foundation (grant identifier: R-279-000-512-281) and from Singapore Ministry of Education (a Tier-2 grant; grant identifier: R-279-000-594-112). Smaranika Panda was supported by Singapore Ministry of Education through a PhD scholarship. RM and MF also thank Natural Science and Engineering Research Council for funding. The illustration in **Fig. 7** was created using BioRender and Adobe illustrator.

## Author contributions

S.P., K.Z. and R.M. conceived the study. S.P., K.Z., R.M., M.F., and E.H. designed the experiments. S.P. executed most of the experiments and collected data. S.P., K.Z., R.M., M.F., and E.H. analyzed the data. J.Z. contributed to gene inactivation. X.M. contributed to plasmid construction. V.F. contributed to the transcriptome analysis. S.P. wrote the original version of the manuscript. S.P., K.Z. and R.M. edited the manuscript. All authors have read and approved the final version of this manuscript. The authors declare no conflict of interest.

## Supporting information

**Supplementary Fig. 1-5**, **Supplementary Table 1-2,** and **Supplementary Note S1** are available in the Supplementary Information file.

