## Supplementary information for "Engineering *Escherichia coli* to produce aromatic chemicals from ethylene glycol"

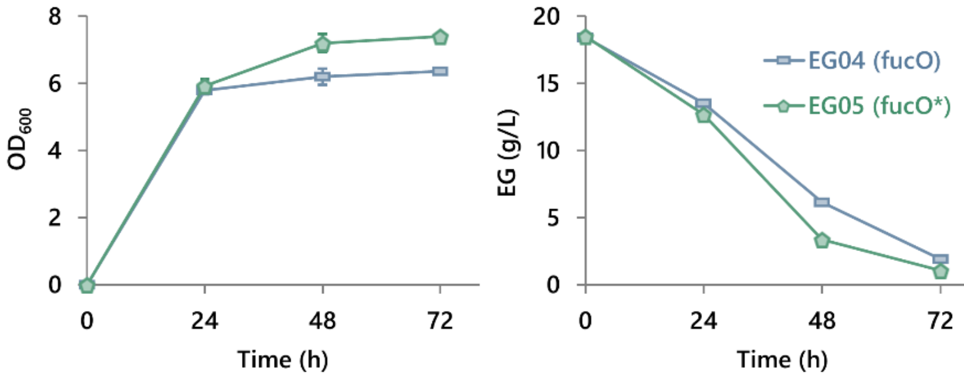


**Supplementary Fig. 1** Effect of mutating fucO (mutations: I7L, L8V to improve the enzyme's oxygen tolerance) on cell growth and EG utilization. **EG04** harbours the plasmid pEG03 (expressing *fucO* and *aldA* under a constitutive promoter). **EG05** harbours the plasmid pEG04 (*fucO* was replaced by *fucO**). **EG04** and **EG05** were grown in a chemically defined medium (10 mL of culture in 125 mL shake flask) containing 10 g/L EG and 1.6 g/L complete supplement mixture (CSM). No antibiotic was added in the medium. The growth temperature was 30 °C. Error bar indicates standard error (n=3).

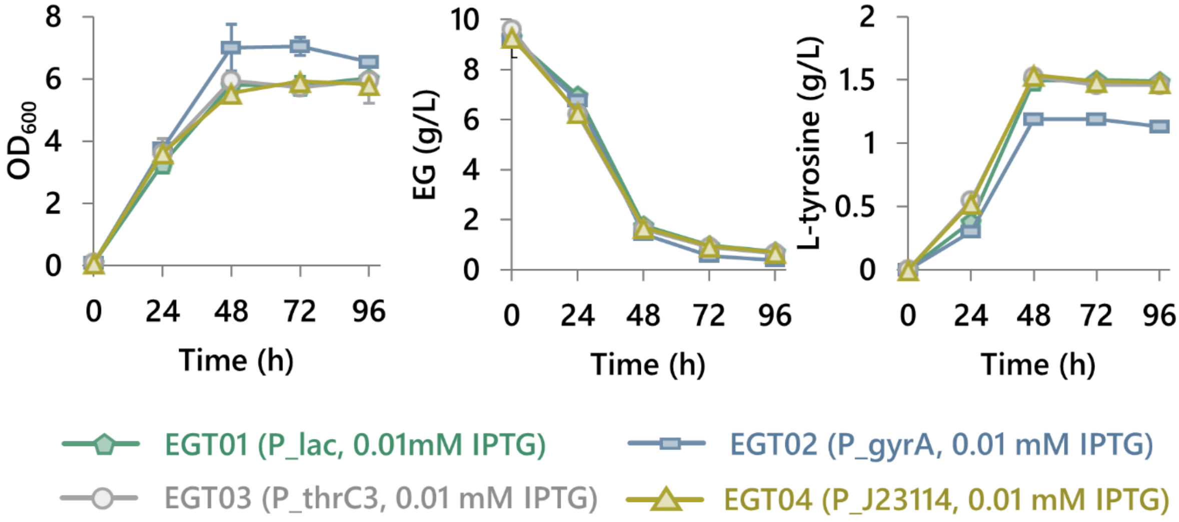


**Supplementary Fig. 2** Effect of promoters to drive *aroG** and *tyrA** on L-tyrosine production. The genotype of the strains **EGT01**, **EGT02**, **EGT03** and **EGT04** can be found in **Table 1**. All the strains were grown in a chemically defined medium (10 mL of culture in 50 mL tube) containing 10 g/L EG and 1.6 g/L CSM. The growth temperature was 30 °C. Error bar indicates standard error (n=3).

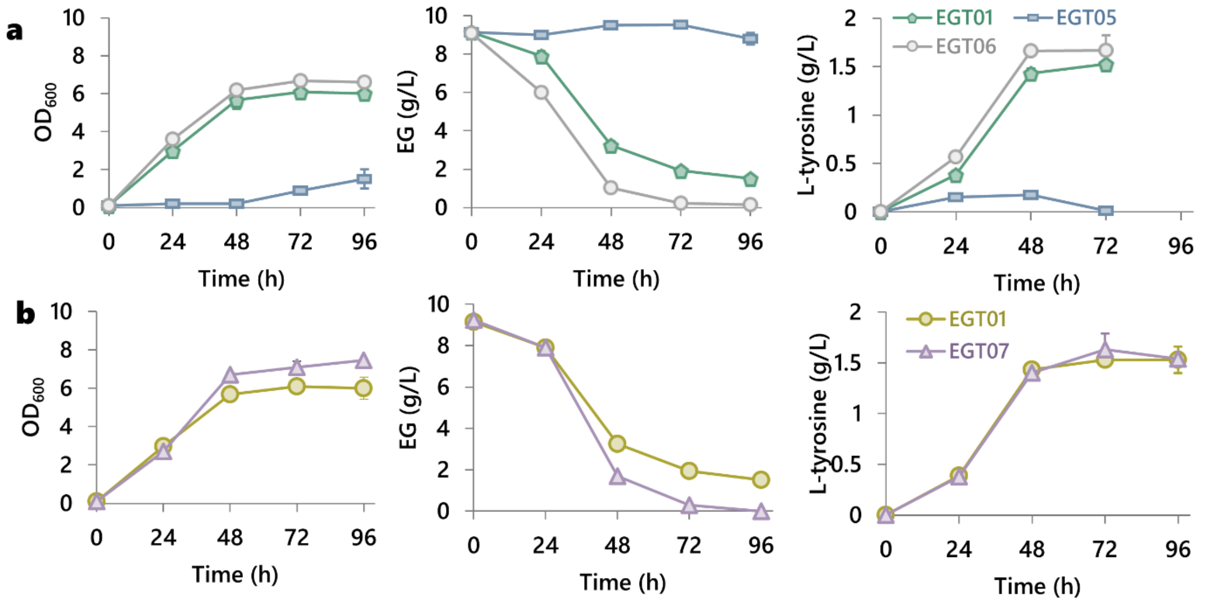


**Supplementary Fig. 3** Optimization of gene expression to further improve L-tyrosine titer. (**a**) Effect of promoters to drive *fucO** and *aldA* on L-tyrosine production. (**b**) Effect of weak constitutive promoter to drive fucO*, aldA, and tyrA*, aroG*. The genotype of the strains **EGT01**, **EGT05**, **EGT06** and **EGT07** can be found in **Table 1**. All the strains were grown in a chemically defined medium (10 mL of culture in 125 mL shake flask) containing 10 g/L EG and 1.6 g/L CSM. The growth temperature was 30 °C. Error bar indicates standard error (n=3).

**Supplementary Note S1**: To test the hypothesis that L-tyrosine production was growth-associated, we conducted two experiments. After the cells were cultured in Medium 1 for 48 h, half of the culture was transferred to a new vessel and mixed with an equal volume of fresh Medium 1. The new culture contained approximately 6.2 g/L EG. The cells grew and converted most of the EG into 1.5 g/L tyrosine in the next 48 h (**Supplementary Fig. 4a**), confirming that the cells in the stationary phase could be activated again to produce L-tyrosine when the cells grew again. But this activation was not achieved if the same cells were removed from the growth medium and resuspended in Phosphate Buffered Saline (PBS) supplemented with 10 g/L EG (**Supplementary Fig. 4b**). The cells could not grow in the PBS solution, because there was no nitrogen source. The results of the two experiments supported that L-tyrosine production was growth-associated and motivated us to improve cell growth through medium optimization.

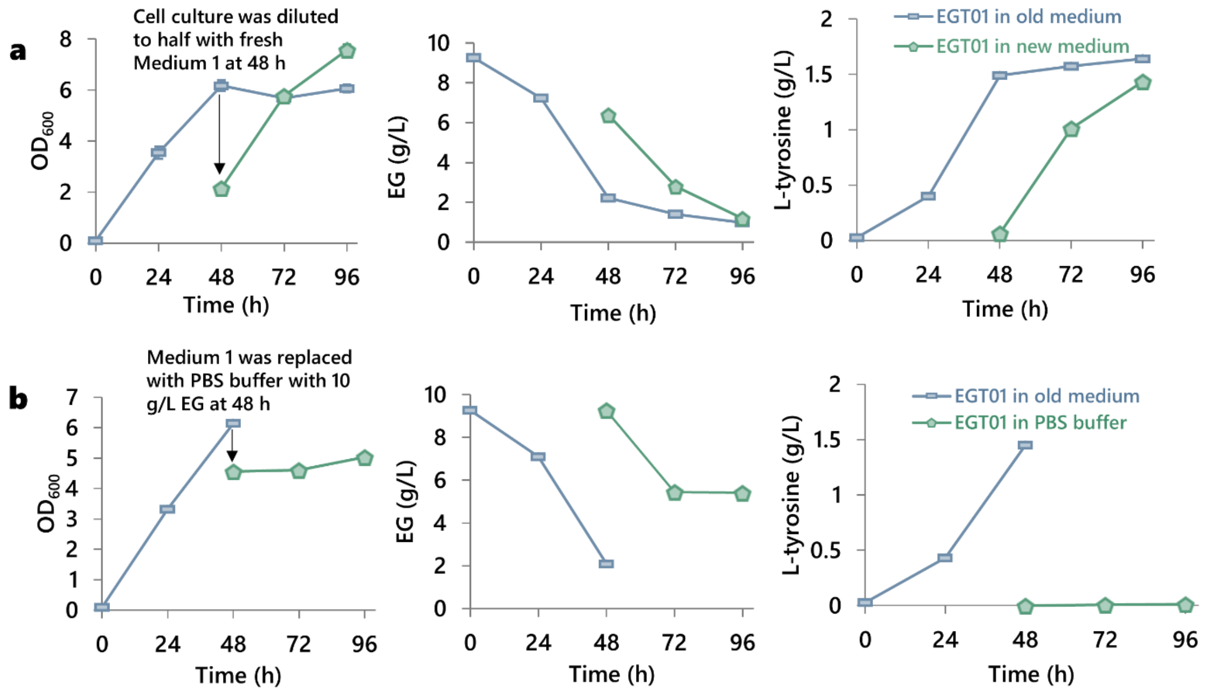


**Supplementary Fig. 4** L-tyrosine production was growth associated. (**a**) Cells in the stationary phase could be activated to produce L-tyrosine when the culture was diluted with fresh medium. (**b**) Resting cells could not convert EG into L-tyrosine, supporting that the L-tyrosine production was growth-associated. Cells in the stationary phase were resuspended in 1X PBS buffer with 10 g/L EG. In all the experiments, *E. coli* **EGT01** was grown in Medium 1 (Materials and Methods) until the dilution or medium replacement. The cells were induced with 0.01 mM IPTG upon inoculation of Medium 1. The temperature was 30 °C in all the experiments. 10 mL culture was grown in 50 mL tube. Error bar indicates standard error (n=3).

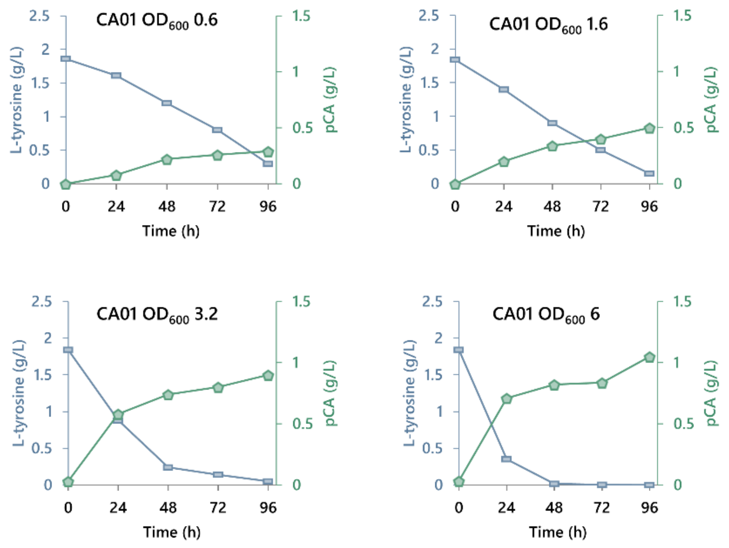


**Supplementary Fig. 5** Conversion of L-tyrosine into pCA by a Tal-expressing strain (**CA01**) at various cell concentrations. In the first stage, **EGT08** was cultured in Medium 1 (10 g/L EG and 1.6 g/L CSM, 0.01 mM IPTG upon inoculation, 10 mL culture in 125 mL shake-flask) for 96 h. **EGT08** was removed by centrifugation. The spent medium containing 2 g/L L-tyrosine produced in stage one was not disturbed. In the second stage, the spent medium obtained from the first stage was used to resuspend **CA01** at different concentrations ($OD_{600}$ 0.6 to $OD_{600}$ 6). The resuspended cells were incubated at 30 °C for 96 h. The **CA01** cells had been cultured separately in LB medium for Tal expression. Error bar indicates standard error (n=3).

**Supplementary Table 1** Primers synthesized in this study.

| Genes over-expressed | Primer orientation | Sequence |
|---|---|---|
| *fucO** | Forward | G**gctaacagaatgctggttaacgaaacggcatggtttgg* |
| | Reverse | A*ccaggcggtatggtaaagct |
| *aldA* | Forward | G*tcagtacccgttcaacatcct |
| | Reverse | A*agactgtaaataaaccacctgg |
| *tyrA** | Forward | G*gttgctgaattgaccgca |
| | Reverse | A*ctggcgattgtcattcgc |
| *aroG** | Forward | G*aattatcagaacgacgatttacgcatcaaa |
| | Reverse | A*cccgcgacgcgctttta |
| *pheA** | Forward | G*acatcggaaaacccgttact |
| | Reverse | A*ggttggatcaacaggcactacg |
| *tal* | Forward | G*acccaggttgttgaacgt |
| | Reverse | A*gccaaaatctttaccatctgc |

**Supplementary Table 2** Mediums and their compositions used in the study.

| **Medium** | **Carbon source; Concentration** | **$NH_4^+$ concentration** | **CSM concentration** |
|---|---|---|---|
| **1** | EG; 10 g/L | 1 g/L | 1.6 g/L |
| **2** | EG; 20 g/L | 1 g/L | 1.6 g/L |
| **3** | EG; 10 g/L | 2 g/L | 1.6 g/L |
| **4** | EG; 20 g/L | 2 g/L | 1.6 g/L |
| **5** | Glucose; 10 g/L | 2 g/L | 1.6 g/L |
